# Predicting interchain contacts for homodimeric and homomultimeric protein complexes using multiple sequence alignments of monomers and deep learning

**DOI:** 10.1101/2020.11.09.373878

**Authors:** Farhan Quadir, Raj Roy, Randal Halfmann, Jianlin Cheng

## Abstract

Deep learning methods that achieved great success in predicting *intrachain* residue-residue contacts have been applied to predict *interchain* contacts between proteins. However, these methods require multiple sequence alignments (MSAs) of a pair of interacting proteins (dimers) as input, which are often difficult to obtain because there are not many known protein complexes available to generate MSAs of sufficient depth for a pair of proteins. In recognizing that multiple sequence alignments of a monomer that forms homomultimers contain the co-evolutionary signals of both intrachain and interchain residue pairs in contact, we applied DNCON2 (a deep learning-based protein intrachain residue-residue contact predictor) to predict both intrachain and interchain contacts for homomultimers using multiple sequence alignment (MSA) and other co-evolutionary features of a single monomer followed by discrimination of interchain and intrachain contacts according to the tertiary structure of the monomer. Allowing true-positive predictions within two residue shifts, the best average precision was obtained for the Top-L/10 predictions of DNCON2: 22.9% for homodimers, and 17.0% for higher order homomultimers. In some instances, especially where interchain contact densities are high, the approach predicted interchain contacts with 100% precision. We show that the predicted contacts can be used to accurately construct the structure of some complexes. Our experiment demonstrates that monomeric multiple sequence alignments can be used with deep learning to predict interchain contacts of homomeric proteins.

## Introduction

Proteins are one of the most important and heavily studied biological molecules. While most proteins form individual three-dimensional structures, they tend to interact with each other to gain functional properties. In fact, most proteins are symmetrical oligomeric complexes with two or more subunits (Goodsell & Olson, 2000), and approximately two-thirds of human enzymes are homo-oligomers (Matthews, 2012).

Since the functionality of most proteins heavily depends on homo-oligomerization, and wet laboratory experiments with actual proteins are time-consuming and expensive, there is a great need to develop accurate computational tools to make such predictions quickly.

Machine learning methods have been developed to facilitate computational modeling of both protein tertiary structures and quaternary structures. However, most of the recent focus has been on the development of computational tools for predicting intrachain (within the same chain) residue-residue contacts and distances to guide tertiary structure modeling (Hopf et al., 2014; Zhou et al., 2018). Some of these methods have performed well in the 12^th^ and 13^th^ Critical Assessment of Techniques for Protein Structure Prediction (CASP) competitions (Adhikari et al., 2018; Alquraishi & Valencia, 2019; Cheng et al., 2019; J. Hou et al., 2020; Schaarschmidt et al., 2018; Senior et al., 2020; Shrestha, Fajardo, Gil, Fidelis, Kryshtafovych, Bohdan Monastyrskyy, et al., 2019; Shrestha, Fajardo, Gil, Fidelis, Kryshtafovych, Monastyrskyy, et al., 2019; Wang et al., 2017, 2018; Xu & Wang, 2019).

Unlike tertiary structure modeling, most of the tools on protein complexes are developed to classify whether two proteins are in contact or not, with some tools developed exclusively for docking tertiary structures together. Only a small number of tools were developed to predict interchain contacts leveraging interchain residue-residue co-evolutionary signals embedded in the multiple sequence alignments of a pair of interacting proteins (i.e., interlog) (Hopf et al., 2014; Ovchinnikov et al., 2014; Zhou et al., 2018). Relevant to the present work (Zhou et al., 2018) and (Zeng et al., 2018) predicted interchain contacts using a deep learning-based intrachain contact prediction tool (RaptorX-ComplexContact) without training the system for interchain contact prediction. Their work involved generating MSAs using homology-based, phylogeny-based, and genome-based interlog (interacting homologs of protein complexes) searches, which is similar to methods employed by (Ovchinnikov et al., 2014) and (Hopf et al., 2014) to generate MSA. Baker (Ovchinnikov et al., 2014) used a pseudo-likelihood-based covariance approach to predict the interprotein contacts in bacterial proteins. The method involved the computation of residue-residue coupling strength between all the interacting protein pairs in the MSA based on the GREMLIN model. The coupling strengths were then ranked and used to compute a score, which was used to derive the distance restraints. EVcomplex (Hopf et al., 2014) used evolutionary couplings (EC) to predict the interface contacts between prokaryotic proteins. Applying EVcouplings (Hopf et al., 2019) to the paired MSA and using the pseudo-likelihood maximization (PLM) approach, both inter-EC and intra-EC were obtained. The normalized raw reliability score (EVcomplex score) was calculated using the inter-EC portion. The interchain residue pairs were ranked according to the EVcomplex scores, and the pairs with scores beyond 0.8 were considered to be in contact with high confidence.

The works above focused on determining the *heteromeric* interprotein contacts and require MSAs of homologous interlogs as input. However, obtaining MSAs of interlogs of sufficient depth is very hard for most proteins because there are many fewer known protein complexes than known monomers. However, the situation can be much different for homo-oligomers because their units are the same monomer. Therefore, both intrachain and interchain residue-residue co-evolutionary signals exist within the same multiple sequence alignment of the monomer. That is, both the co-evolution signal between residue i and residue j within a monomer and that between residue i in the monomer and residue j in its identical partner are mixed in the same multiple sequence alignment of the monomer. This phenomenon has been recognized before but has not been leveraged to predict interchain contacts in homo-oligomers (Uguzzoni et al., 2017).

The objective of this study focuses on predicting the interprotein contact of *homomeric* proteins (chains having similar amino acid sequences) using MSAs of the monomer unit in homo-oligomers. Similar to (Zeng et al., 2018; Zhou et al., 2018), our approach can be applied to both prokaryotic and eukaryotic proteins wherever sufficiently deep MSAs are available. Unlike the above approach to generate MSAs, ours directly extracts co-evolutionary features of homomeric proteins from homology-based MSAs constructed without any special genome- and phylogeny-based methods. We then directly apply DNCON2 (Adhikari et al., 2018)), a deep learning method trained to predict intrachain contacts, to these features to predict both intrachain and interchain contacts, which we distinguish according to the tertiary structures of monomers (See Supplementary Section Figure S1 for the DNCON2 deep learning architecture, and Table S1 for the list of features used by DNCON2). This paper focuses on proving this concept, while future work will focus on developing an exclusive deep-learning-based interprotein contact predictor.

Interchain protein contact predictions are valuable not only to identify protein-protein interactions, but also to inform the construction of the proteins’ 3D complex structures. Some docking tools use interchain contacts and distances as additional restraints to compute better quaternary structures. Since more accurate interprotein contacts lead to better quality quaternary structures, this research will pave the way to improve *de novo* interprotein contact prediction and complex structure construction methods.

## Methods

### Dataset preparation

The lists of homo-oligomers were obtained from the 3DComplex dataset (Levy et al., 2006). We selected our protein dataset from this massive pool of oligomeric proteins after the Protein Data Bank (PDB) files underwent a cascade of filtration steps. The first of our filters selected from this list only those proteins which were homo-oligomers. We then discarded those proteins that had interactions with nucleic acids. The next step cleaned and selected PDB files according to the process described in **Figure 1**.

**Figure 1:**
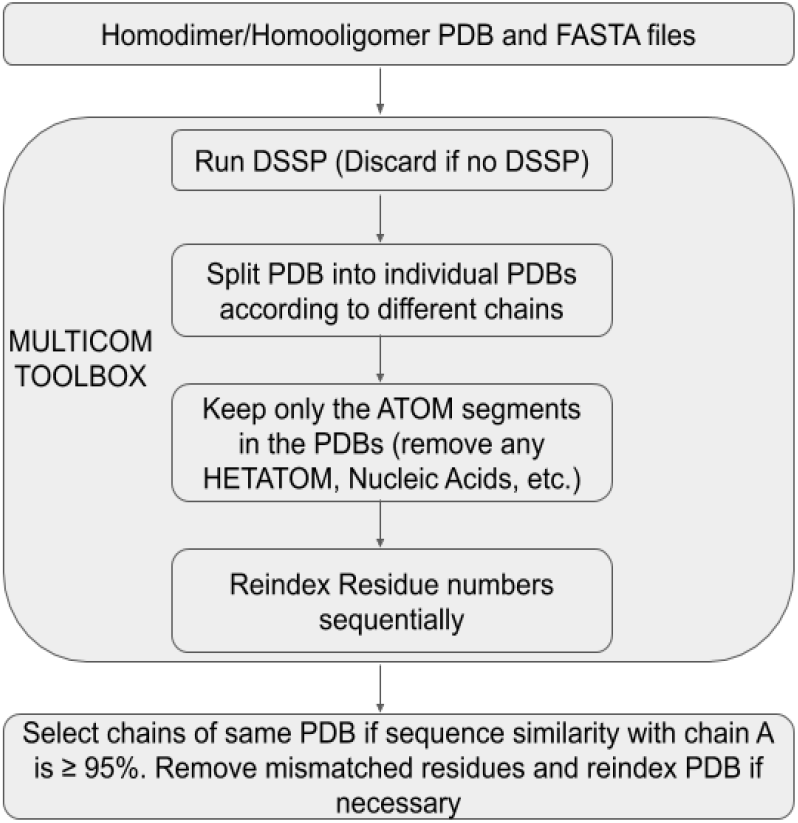
Diagram describing how the input PDB file was pre-processed using the MULTICOM TOOLBOX to clean up the PDB files. If no DSSP is available, the PDB was removed from our list. The individual chains in the multimer PDB were separated into individual files containing the ATOM (x, y, and z coordinates) segments only while discarding all other information. Only chain pairs whose FASTA sequences match 95% or more were kept, and any mismatched residues were removed to ensure homogeneity between chain pairs. The residue numbers were reindexed to ensure continuity, thereby removing any chain breaks and negative residue numbers.

Since Protein Data Bank (PDB) files tend to contain chain breaks and more information than necessary, we cleaned them using MULTICOM Toolbox (Cheng et al., 2012; Jie Hou et al., 2020) as shown in Figure 1. It applies DSSP (Kabsch & Sander, 1983) to generate secondary structure and solvent accessibility information for each PDB file. The toolbox first checks if DSSP output can be generated for a protein and, if so, splits the entire PDB file into individual files corresponding to the chains present, keeping only the ATOM (x, y, and z coordinate) portion. The residue numbers were then reindexed to ensure every residue number begins with a ‘1’ and continues without any breaks. We then checked to see if the FASTA sequence between all chains is similar or not. We discarded any pairs of chains if the pairwise sequence similarity is less than 95%. The individual chain-wise PDB files were further processed to remove the mismatched residue information, and residue numbers were reindexed if necessary, for homogenizing the residue similarity between chains.

This list of cleaned protein chains was then used to remove redundant sequences with more than 30% sequence identity using the software “mmseqs” (Hauser et al., 2016). We excluded the proteins which did not have any interchain contact (distance between interchain residues was > 6.0 Å). Several proteins that failed with FreeContact (Kaján et al., 2014), PSICOV (Jones et al., 2012), and PSI-BLAST (Altschul et al., 1997) in the feature generation process of DNCON2 were also excluded. Finally, we obtained a dataset of 8681 homodimeric proteins and 6764 higher order homo-oligomeric (hereafter, “homomultimeric”) proteins for prediction and analysis. The homodimeric and homomultimeric datasets did not overlap and were treated separately.

### Prediction and Evaluation of interchain contacts

Since dimeric proteins contain two chains, the underlying process is relatively simple. Hence, in this study, we only describe predicting the homomultimeric proteins’ interchain contacts, which implicitly covers the dimers. As described in **Figure 2**, the cleaned chain-wise split homomultimer PDBs and FASTA sequences were given input to our prediction and evaluation system. The coordinates of chain ‘A’ (the first chain) were used to calculate the intrachain residue-residue distance. An *intrachain* contact between two residues i and j is said to exist if the Euclidean distance between the respective C_ (C_ for glycine) atoms of residues i and j is less than or equal to 8.0 Å according to the definition used in DNCON2 (Adhikari et al., 2018; Jie Hou et al., 2020). If the C_ was not found in the PDB file, we chose C_ instead. We obtained the true intrachain contact map using this rule.

**Figure 2:**
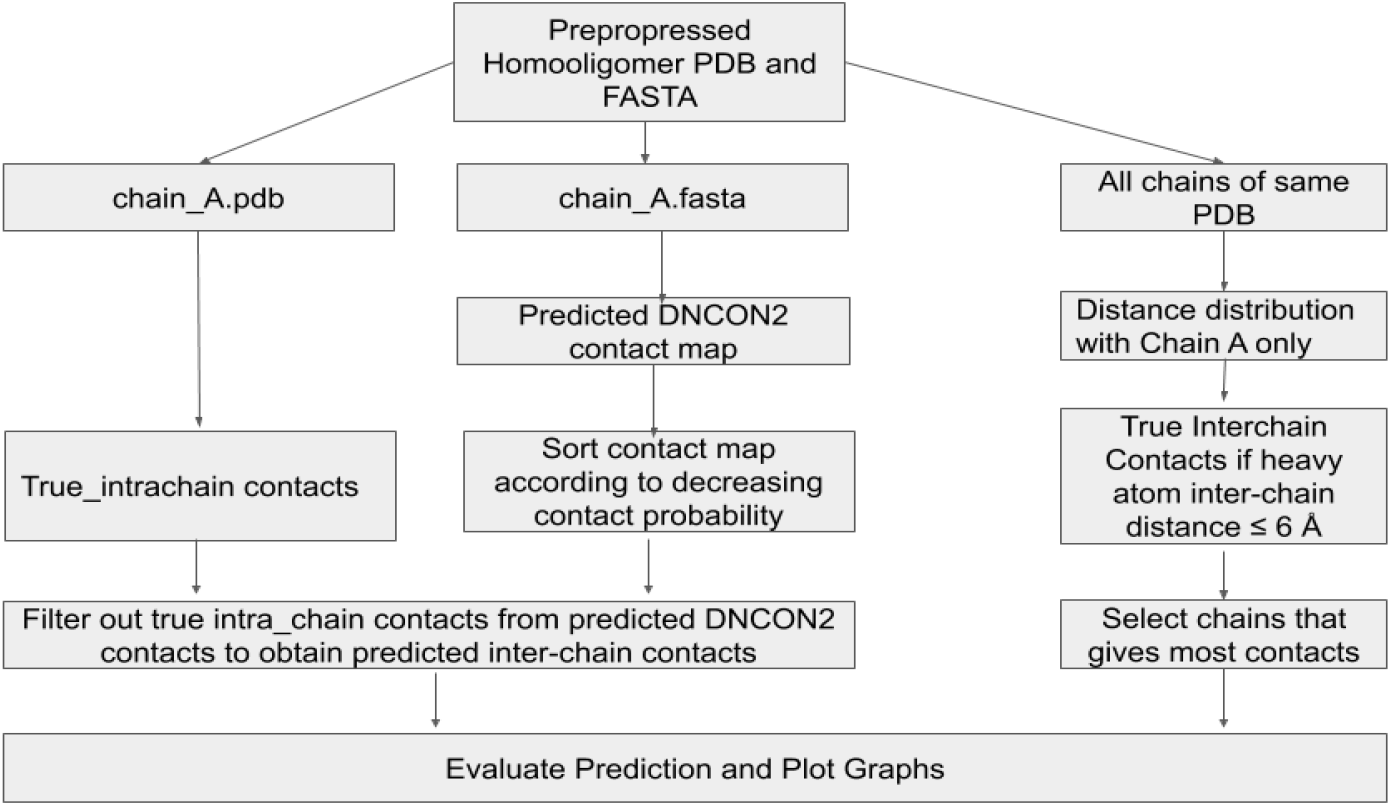
Workflow diagram describing how the pre-processed input PDB and FASTA sequence was used to derive true intrachain contacts, true interchain contacts, predicted interchain contacts, and finally, obtain the evaluation and visualization of the prediction.

We also defined *interchain* contact between chains in a protein if the Euclidean distance between the heavy atoms of the residues in the respective chains is less than or equal to 6.0 Å (Hopf et al., 2014; Ovchinnikov et al., 2014; Zhou et al., 2018). We obtained a pairwise contact list between the first chain (chain A) and the other chains after separating the homomultimeric PDB into separate chains. For example, if a protein contains four chains A, B, C, and D, we pair them as AB, AC, and AD and determine the contacts between these pairs according to the definition described above. From this distribution, we selected the pair with the highest number of contacts and based the rest of our analysis on this pair as we compare the true interchain contacts with the predicted interchain contacts to obtain our predictor’s precision.

To predict the interchain contacts of a protein chain, the FASTA sequence of the first chain (i.e., chain A) was fed into our predictor program-DNCON2 - which outputs the predicted intrachain contact map. This predicted contact map was then processed to filter out the short-range contacts (contacts whose sequence separation is less than 6, i.e., for contact positioned at i and j, if |i-j| < 6, we discard this contact). We also reported the precision of the intrachain prediction in our results. This short-range removed intrachain contact map was post-processed as follows to obtain the interchain contacts:

1. We removed the matching true intrachain contacts from the prediction. Our assumption states that the remaining contacts in the prediction contact map should correspond to some sort of interchain contact.
2. We then computed the precision using the contact map file obtained from (1) and the true interchain contacts using InterConEva- an extension of ConEva (Adhikari et al., 2016) tool for interchain contact evaluation. It should be noted that DNCON2 outputs the upper triangle of the prediction matrix since the intrachain contact map is symmetric. However, in interchain prediction, the entire matrix needs to be considered since contact between residues i and j of two different chains is distinctly different from contact between residue j and i. Therefore, if the final predicted contact map had a contact (x, y), we checked for both (x, y) and (y, x) in the true interchain contact map and considered them to be two separate contacts.

We experimented further by applying relaxations to the above parts (1) and (2). We termed them as “relax removal,” and “ relaxation,” respectively. During relax removal, we removed the true intrachain contacts from the predicted contact map as follows:

1. If position (i,j) is a true intrachain contact, and the relax parameter is n (where, n=0, 1, and 2), then let X = [i-n,i+n] and Y= [j-n, j+n]
2. Remove all contacts (X_p_,Y_q_) from the predicted contact map where X_p_={i-n, i-n+1, …, i+n} and Y_q_={j-n, j-n+1, …, j+n}. This removes (sets to zero) a square matrix of dimension n x n centered at (i,j) from the predicted intrachain contact map.

Relaxation follows a similar approach. If a prediction is found for the position (i,j), we look for contacts within the square matrix of dimension n x n centered at (i,j) in the true binary interchain contact map. If any nonzero value is found within the n x n square region centered at (I,j) of the true interchain contacts, it is counted as a successfully predicted contact. The value of n is ranged from 0 to 2 since we perform relaxation within two residue shifts.

We also selected some best-case results and constructed their complex structure. We compare interchain contact maps using InterConeva, visualize quaternary structures using Chimera (Pettersen et al., 2004), and calculate TMscore similarity scores using TM-Align (Zhang & Skolnick, 2005).

## Results and Discussions

### The Precision of Intrachain Contact Prediction

**Table 1** shows the average precision of intrachain contact predictions for homodimers and homomultimers. Precision is relatively high because both short-range and medium/long-range contacts are considered, and generally good multiple sequence alignments are obtained for the proteins in the dataset. Precision values drop as the number of predicted contacts is increased from Top-5 to Top-2L (L being the protein length). The average intrachain precision for the homomultimers is slightly lower than that of the homodimers.

**Table 1:** Table showing the average precision of predicted intrachain contacts obtained for the homodimers and homomultimers using ConEVA. The sequence separation of contacts is short-range or above (>= 6). L represents the length of the protein sequence.

| Precision (%) |  |  |  |  |  |  |
| --- | --- | --- | --- | --- | --- | --- |
| Dataset | Top-5 | Top-L/10 | Top-L/5 | Top-L/2 | Top-L | Top-2L |
| Dimer | 96.01 | 94.25 | 92.08 | 85.54 | 74.81 | 57.05 |
| Multimer | 89.41 | 87.60 | 85.33 | 79.47 | 70.11 | 54.02 |

### The Precision of Interchain Contact Prediction

The predicted interchain contacts were compared with the true interchain contacts. We have nine scenarios grouped according to nine different combinations of relaxation values (n = 0, 1, and 2) applied to relax removal and relaxation.

The results from **Figure 3** and **Figure 4** show that as we perform more relax removal and relaxation, the interchain precision increases. We obtain a maximum average interchain precision of 22.9% and 17.0% for homodimers and homomultimers, respectively, for the Top-L/10 group with relax removal = 2 and relaxation = 2. When compared with the precision results obtained from random predictions (Supplementary Section Tables S3 and S4), for the Top-L/10 with relax removal = 2 and relaxation = 2, our prediction is 4.1 times greater for homodimers and 3.5 times better than homomultimers. For all groups, increasing relaxation tends to bring about slight increases in precision values. This increase is due to the consideration of true-positive hits over a flexible boundary that enables the system to discover more contacts within a fixed vicinity. There is a more significant increase in precision when we go from relax removal = 0 to relax removal = 1, but slight increase when we move from relax removal = 1 to relax removal = 2. However, from Top-L/5 and beyond, precision drops drastically for all the graphs.

**Figure 3:**
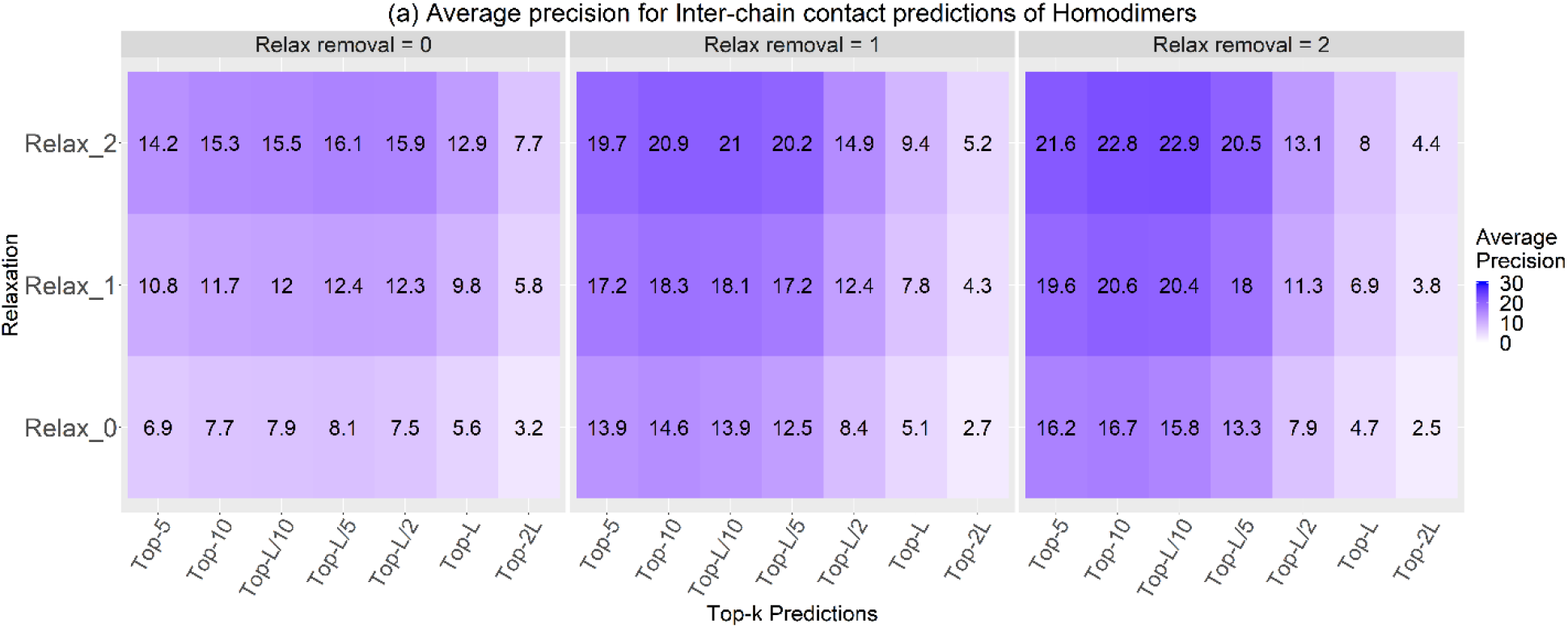
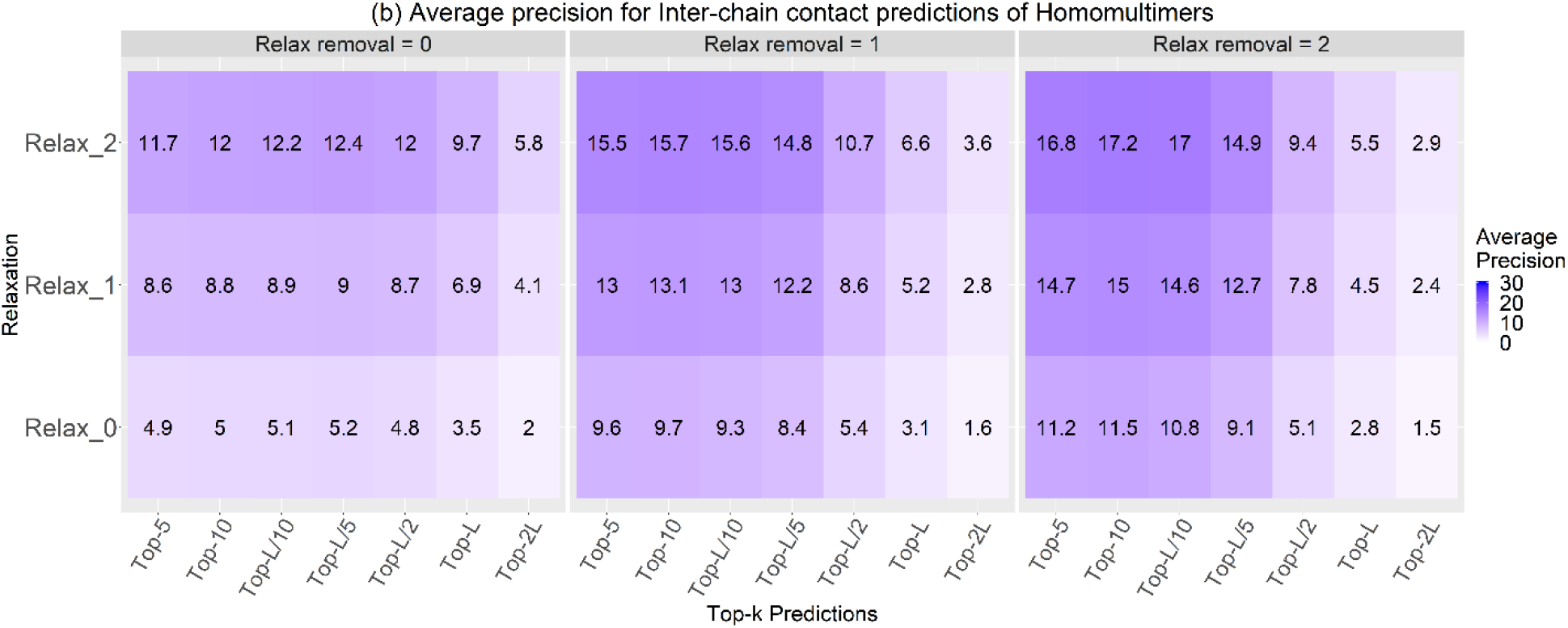
The precision heatmap of interchain contact predictions for the (a) homodimers and (b) homomultimers for the Top-k predictions where k= 5, 10, L/10, L/5, L/2, L, and 2L. For all categories, as we do more relax removal and relaxation, precision values increase within the respective Top-k categories. Relax removal = 2 and relaxation = 2 shows the best precision of 22.9% for homodimers and 17.0% for homomultimers within the Top-L/10 predictions.

**Figure 4:**
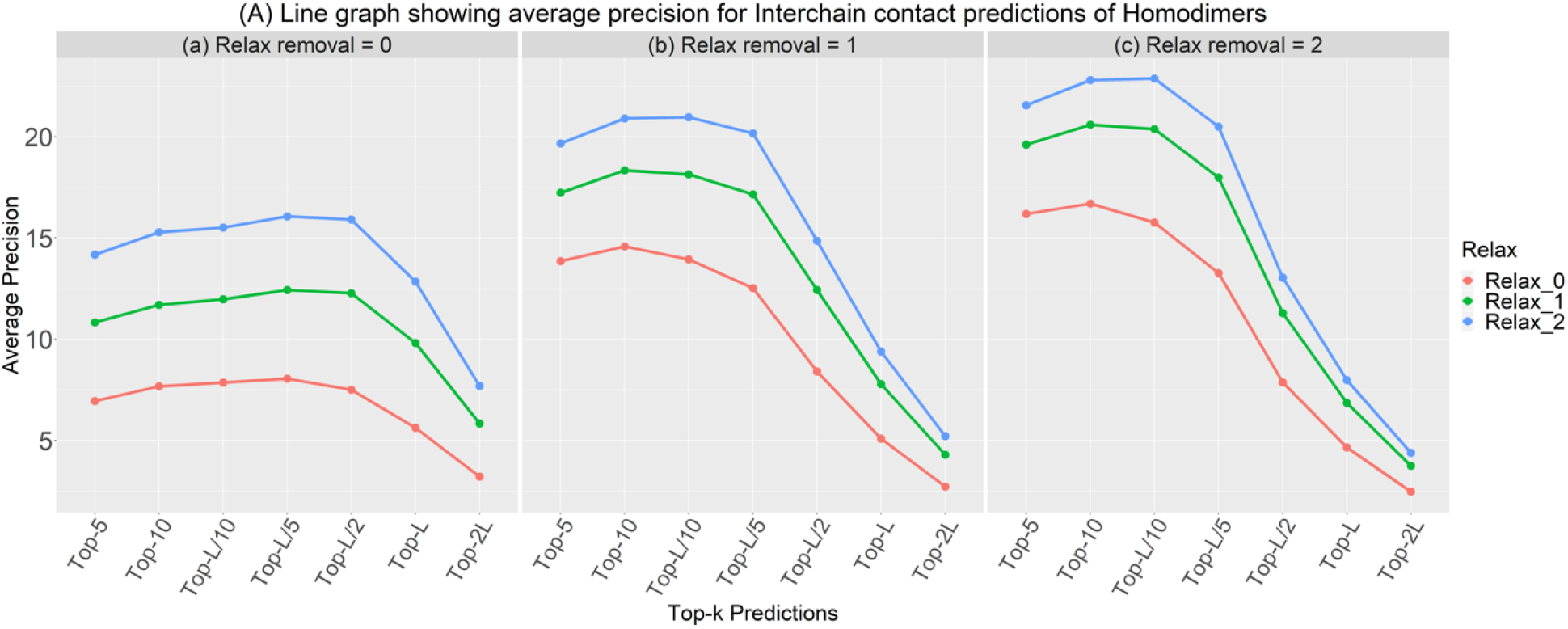
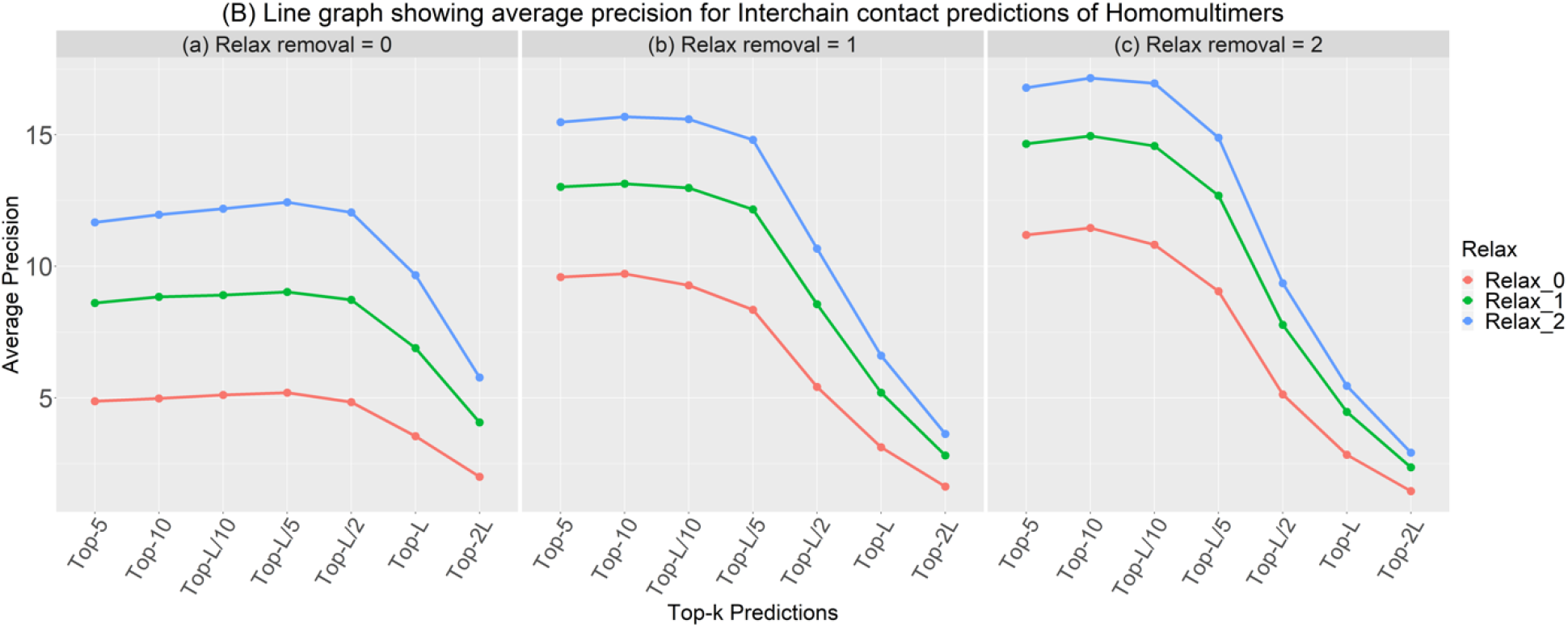
The figure shows how relaxation removal and relaxation affect interchain contact prediction precision (A) for homodimers and (B) for homomultimers. For homodimers, (a) there is no relaxation removal. Precision steadily increases for all relaxation thresholds until Top L/5, then sharply decreases. Similar observations are seen for (b) and (c) with maximum precision occurring at Top-10 for (b) and top-L/10 for (c). Increasing relaxation thresholds from 0 to 2 always increases precision.

We explain this using **Figure 5**, which shows the frequency of proteins (y-axis) for which a given range of contacts (x-axis) was successfully predicted. Relaxation and relax removal were also varied. The number of successful contact predictions is divided into five categories (x-axis), where 0 means no successful contact was predicted for this protein. A protein is tallied in the range 1-20 if the total number of true-positive predictions for this protein is at least one but less than or equal to 20; and so forth. We show only the Top-L/10 (a) and Top-2L (b) graphs for homodimers in Figure 5 in this analysis for simplicity. A bulk of our samples (especially for Top-L/10) remains mispredicted by our predictor (has zero predictions). There are more successful predictions if the number of contacts present in the proteins is within the 1-20 range (Figure 5 (a)). If proteins have more than 20 contacts, the number of successful predictions is low. As we perform relaxation, we can see that the incorrect predictions (zero contact prediction) go down, while successful predictions, especially in the 1-20 range, increases drastically, leading to an increase in precision. As we perform relax removal, the number of proteins in the 1-20 range remains similar, but more well-predicted contacts appear for proteins with more than 20 contacts. This is expected since relax removal removes false-positive contacts from the Top L/10 predictions leading to more true-positive predictions being discovered, thereby increasing precision.

**Figure 5:**
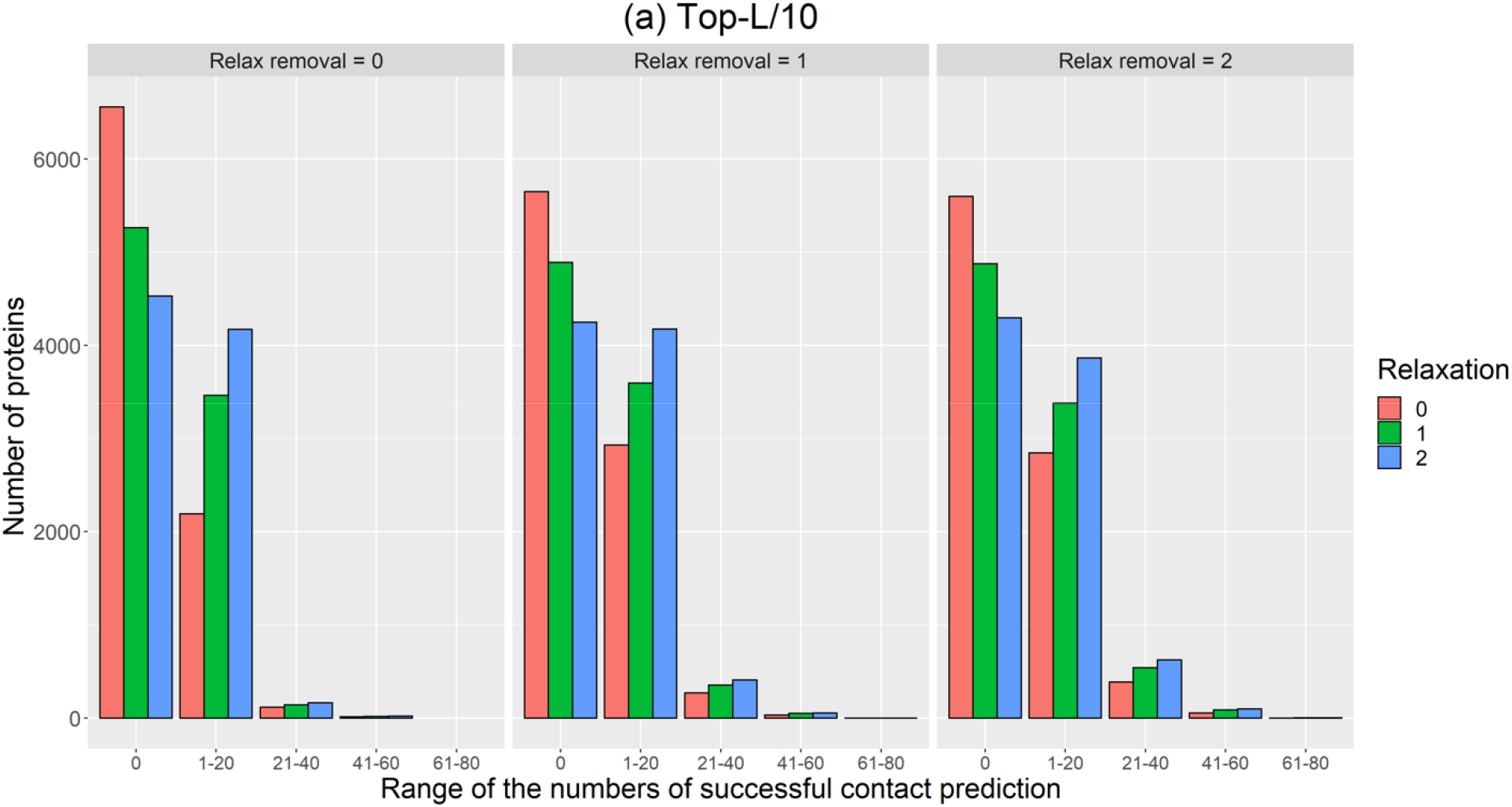
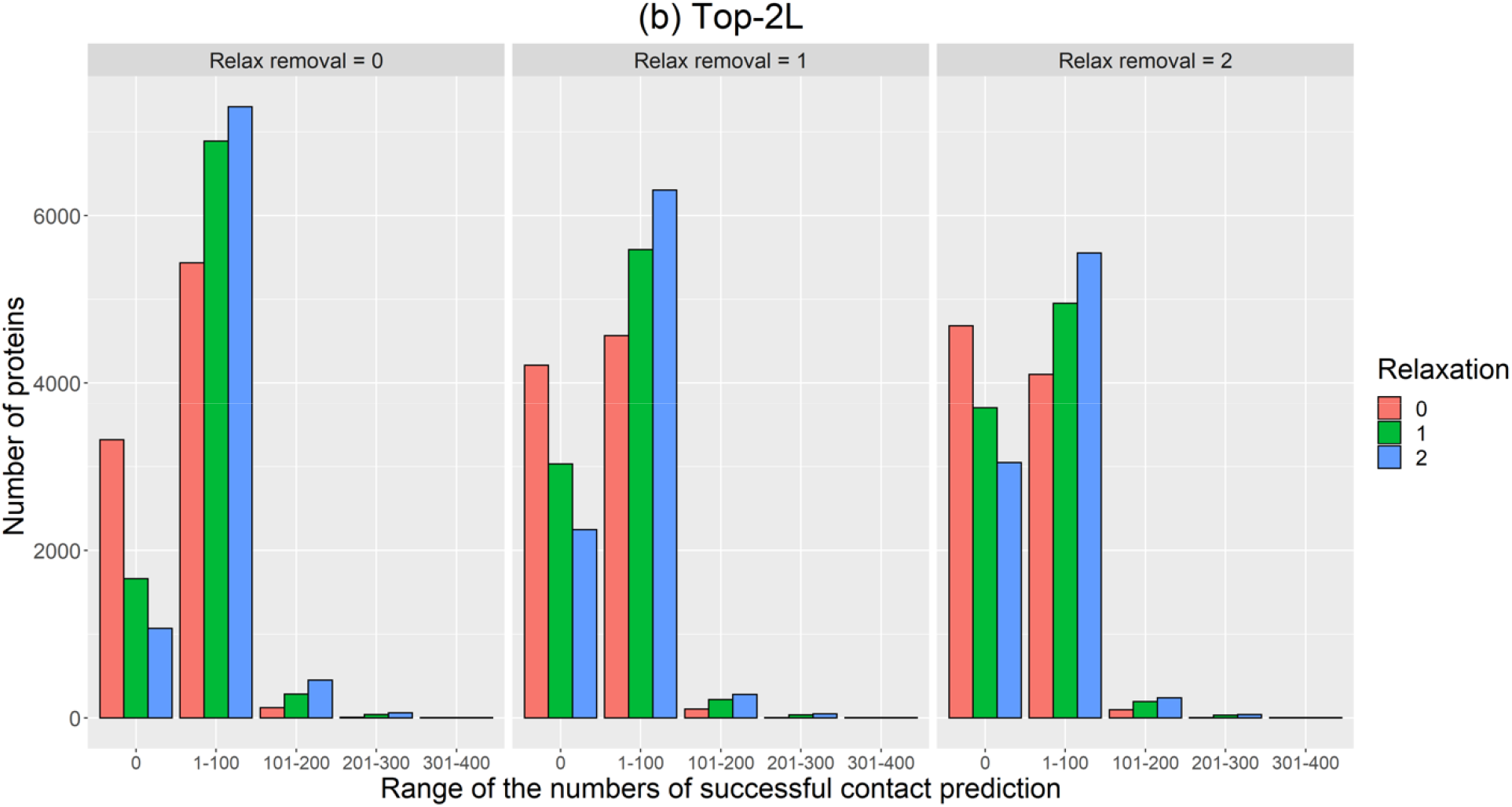
The number of homodimeric proteins (y-axis) for which the total number of contacts were successfully predicted within given ranges of total true-positive contacts (x-axis) at different combinations of relaxations and relax removals for (a) Top-L/10 and (b) Top-2L, respectively. (a) The L/10 shows more mispredictions (zero value in x-axis). Among the proteins whose contacts were successfully predicted, most of the proteins have one to 20 true-positive contacts. (b) The bottom graph shows similar results for Top-2L, but most proteins have total contacts within the 1-100 range. In both graphs, we see relaxation increases the number of proteins with well-predicted contacts. Relax removal for the Top-L/10 group increased the number of proteins having more than 20 true-positive contacts. However, for Top-2L, relax removal decreased the number of well-predicted proteins, especially those with one to 100 contacts.

However, we see sharp precision drops in the cases of Top-L/5 and beyond (Figure 4). We look at the Top-2L graph (Figure 5 (b)) to analyze this observation. Unlike the Top-L/10 graph (Figure 5 (a)), in Top-2L, we see fewer mispredictions (zero value category in x-axis) and more proteins that have been successfully predicted to have 1-100 contacts because 2L is a more expansive range compared to L/10. So more true-positive contacts are encountered. However, during the precision calculation, we are dividing the total number of true-positives by 2L, which is comparatively a higher number. If L is large, the precision drops drastically. As we perform relax removal, we see very little increase in precision for Top-L/5 and beyond, while precision also decreases in some instances (Figure 3 and Figure 4). The Top-2L graph from Figure 5 (b) further suggests that relax removal increases the number of mispredictions (zero value category in the x-axis) while the number of proteins for which successful contacts were predicted decreases. This is due to removing some well-predicted interchain contacts from the predicted contact map when performing the relax removal. In some cases, we discovered that all the predicted contacts get removed, resulting in precisions to become zero.

In further analysis, we compare contact density with the precision of our prediction. Contact density is the total number of native contacts per length of the protein (number of true interchain contacts / L) (Zhou et al., 2018). Again, we discuss the contact density and precision of the Top-2L predictions of homodimers without any relax removal for simplicity. **Figure 6** shows that precision increases with increasing contact density for the Top-2L group. We observe that performing relaxation from 0 to 1 increases the precision for contact densities of all ranges. In this case, precision almost doubles when contact densities are less than 3.50, while, for contact densities beyond 3.50, the precision increases slightly. Performing relaxation from 1 to 2 increases precision but lesser than what is observed when relaxing from 0 to 1. Relaxation from 1 to 2 affects precision values less for contact densities 3.50 and more. Proteins with high contact density tend to get predicted better than proteins with low contact density. It should be noted that most proteins have very sparse interchain contacts and thus have low contact densities. Also, the number of proteins with high contact densities is significantly less (See Supplementary Section Table S2), so these proteins play little role in the overall precision.

**Figure 6:**
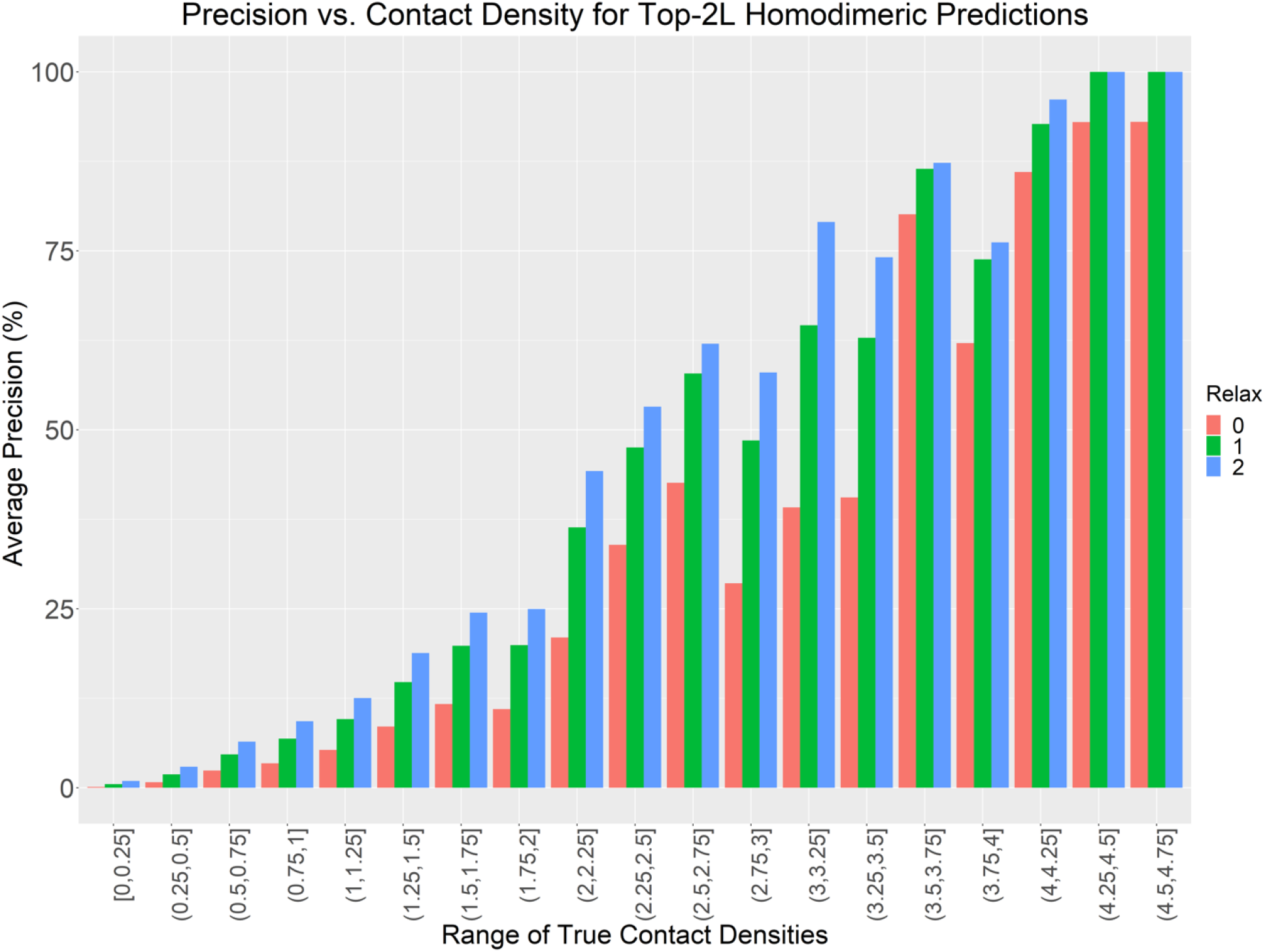
Bar plot depicting the prediction of Top 2L interchain contact predictions changes as contact density varies with no relaxation removal. We can see that high contact density leads to high precision. Relaxation has little effect on precision when contact densities are beyond 3.50.

### Case study of two best predictions

We investigate the results of two best predictions for two proteins (PDB code: **1A64** and **1IHR**) (Tables 3 and 4, respectively)

**Table 2** shows that although DNCON2 predicted intrachain contacts for this target with relatively low precision, all the interchain contacts were correctly predicted, which is also confirmed visually by the bulk overlapping of the green and red spots, as seen in **Figure 7**. Some scattered green dots that do not overlap with the red ones are mirror images to the blue spots indicating that these are good intrachain predictions. These scattered intrachain predictions disappear as we perform relax removal. Using these contact maps, we use CNS (Crystallography and NMR System) (Brunger, 2007; Brünger et al., 1998) to create quaternary structures and visualize them using Chimera (Pettersen et al., 2004) (**Figure 8**). Finally, we evaluate the resulting structures’ accuracy using TMAlign (Zhang & Skolnick, 2005). Our results indicate very close resemblance to the actual structures with almost perfect TM-scores for the relax removed contacts since intrachain contacts are almost eliminated, leaving behind only true-positive interchain contacts.

**Table 2:** The precision (%) of intrachain and interchain contact predictions for PDB 1A64. The last three rows correspond to interchain precision at different relaxation removal levels.

| Relax removal | Relaxation | Top-5 | Top-10 | Top-L/10 | Top-L/5 | Top-L/2 | Top-L | Top-2L |
| --- | --- | --- | --- | --- | --- | --- | --- | --- |
| Intrachain Precision |  | 0 | 40.0 | 33.33 | 52.63 | 46.81 | 38.30 | 32.45 |
| 0 | 0 | 100 | 100 | 100 | 100 | 100 | 100 | 100 |
| 1 | 0 | 100 | 100 | 100 | 100 | 100 | 100 | 100 |
| 2 | 0 | 100 | 100 | 100 | 100 | 100 | 100 | 100 |

**Figure 7:**
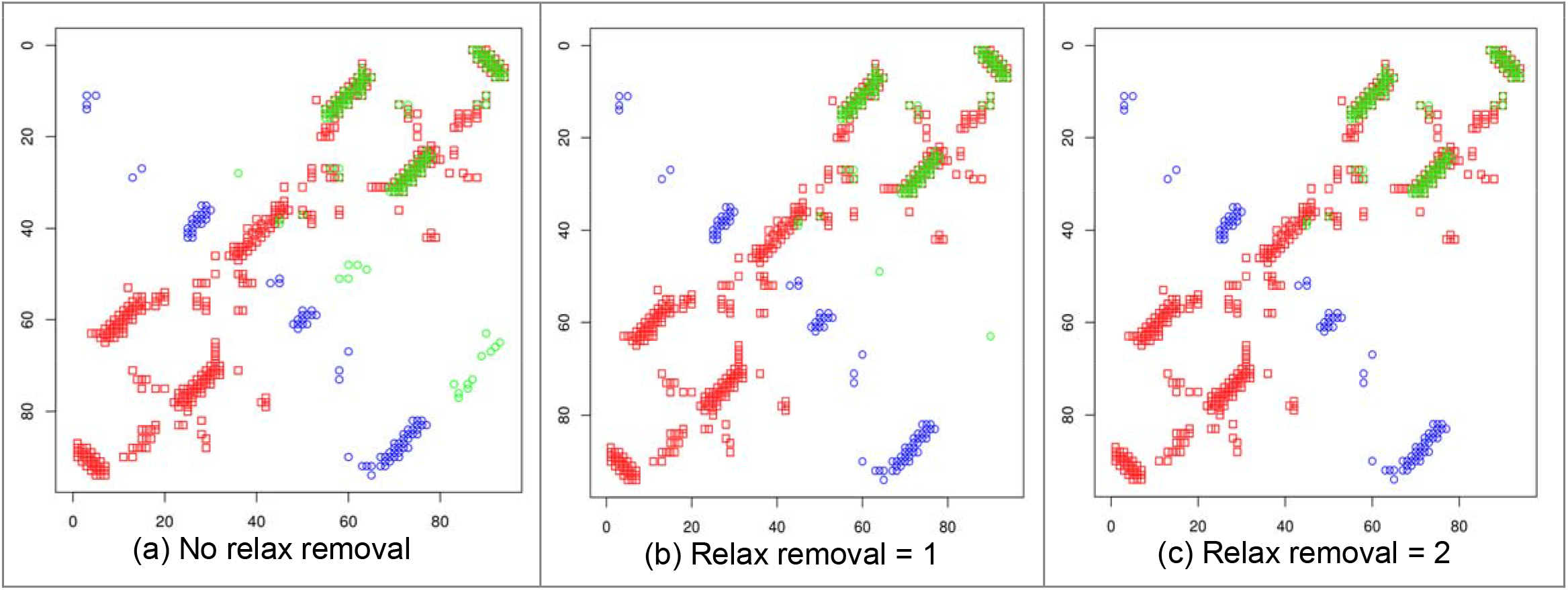
Contact map comparison between true intrachain (blue), predicted interchain (green), and true interchain (red) contacts for 1A64. Since the true intrachain contacts and predicted inter-chain contacts are symmetric, only the lower triangle and upper triangle contacts are, respectively, shown. (a) shows more green spots since no relax removal was done. From (b) to (c), the green contacts become sparse due to removing more predicted contacts assumed to be intrachain. The green dots that overlap with the red dots are correct inter-chain contact predictions.

**Figure 8:**
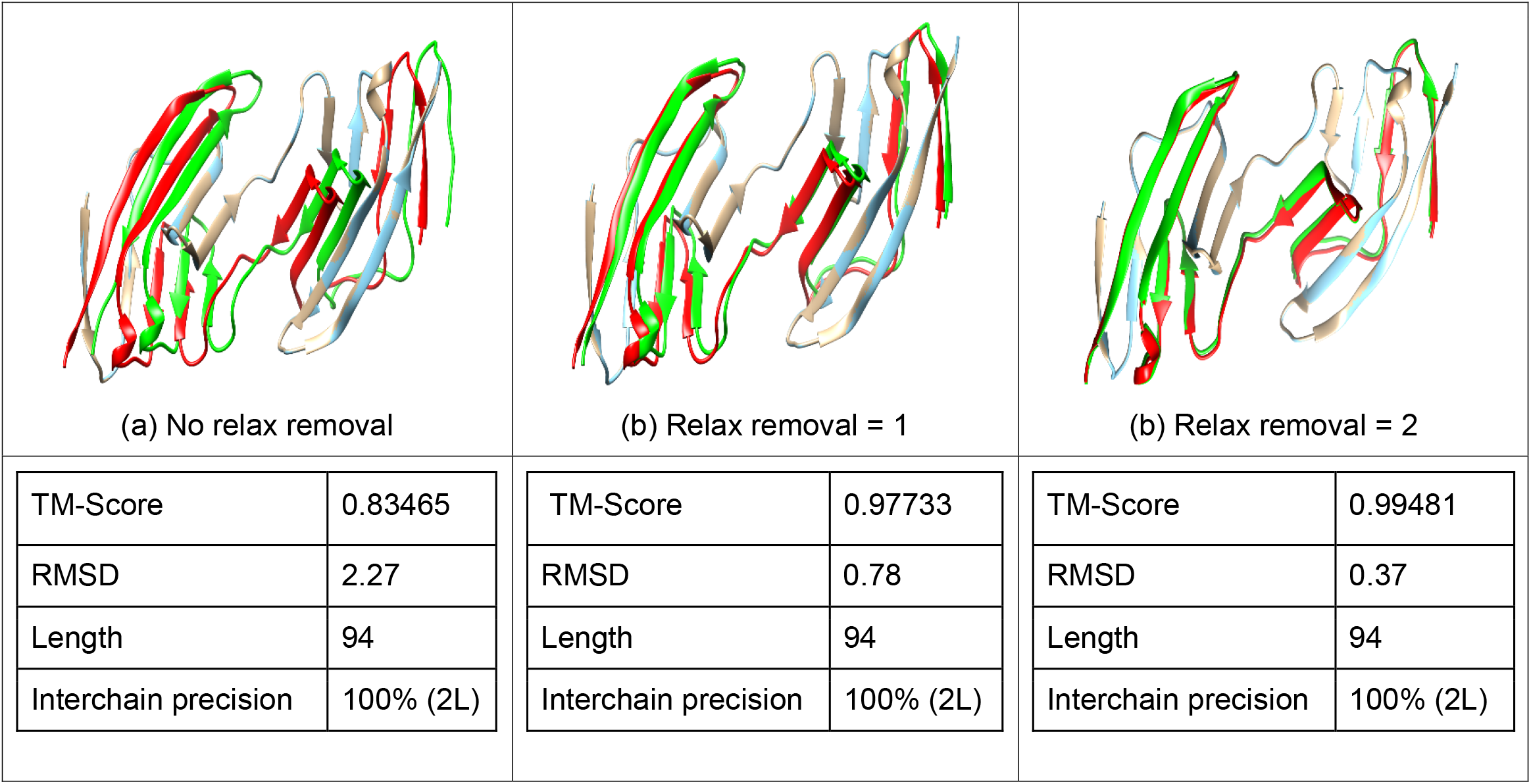
Comparison for target 1A64 between its true homodimer structure and the structure derived from our predicted contacts built by CNS (Crystallography and NMR System). (Golden: original chain A; Cyan: reconstructed chain A; red: original chain B; green: reconstructed chain B) The TM-score and RMSDs were obtained using TM-Align. From (a) to (c), as we perform relax removal, we remove more intrachain contacts to obtain a higher proportion of true-positive interchain contacts (as seen in the previous contact map diagram Figure 7). As a result, the final structures become more accurate, and TM-score increases with decreasing RMSD.

**Table 3** shows the precision values of our predictions for Target 1IHR, which is very similar to observations obtained for 1A64. The contact maps of 1IHR, as visualized in **Figure 9**, also shows large green and red region overlaps indicating near-perfect interchain contact prediction while some scattered predictions correspond to intrachain contacts, which become sparse with relax removal. Structure visualization and TM-score analysis from **Figure 10** shows that as we perform relax removal, the contact map becomes more accurate, resulting in better complex structure formation. Switching from Top-2L to Top-L contacts and applying relaxation removal gives the best TM-score. This can be attributed to the fact that Top-L contacts have 100% precision while Top-2L precision is slightly less. Some structures, like Figure 10(d), give lower TM-score due to a lack of overlap in some of the noodle-like structures and a portion of the α-helix.

**Table 3:**
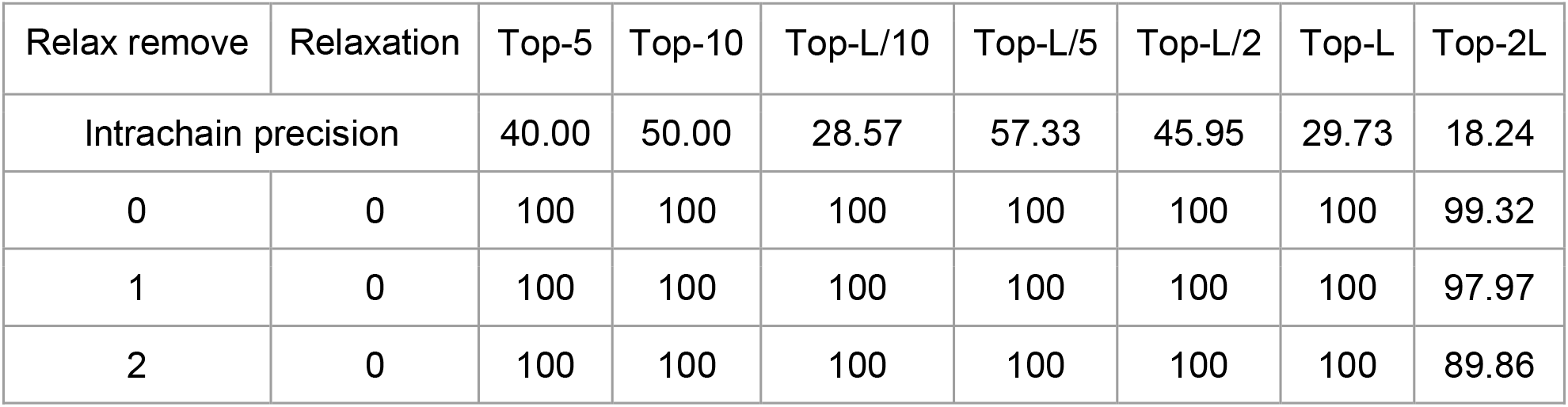
Table showing the precisions (%) of some top predictions done by our system for PDB 1IHR.

**Figure 9:**
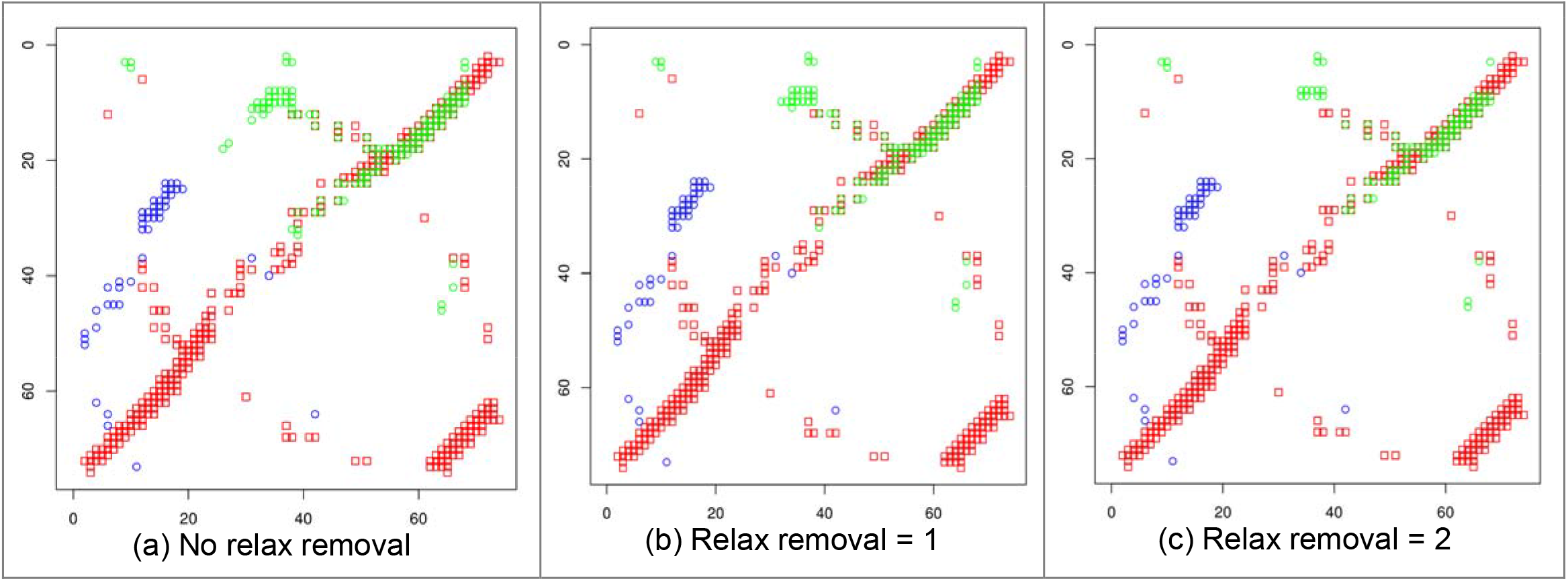
Contact map comparison of the Top-L true intrachain (blue), predicted interchain (green), and true interchain (red) contacts for 1IHR for different relaxation removals. (a) shows more green spots since no relax removal was done. From (b) to (c), the green contacts become sparser due to removing more predicted contacts assumed to be intrachain. According to Table 1, the final Top-5, Top-10, Top-L/10, Top-L/5, Top-L/2, and Top-L precisions for this prediction are all 100%. Only the Top-2L precision drops to 99.32%, 97.97%, and 89.86% for relax removal 0, 1 and 2, respectively.

**Figure 10:**
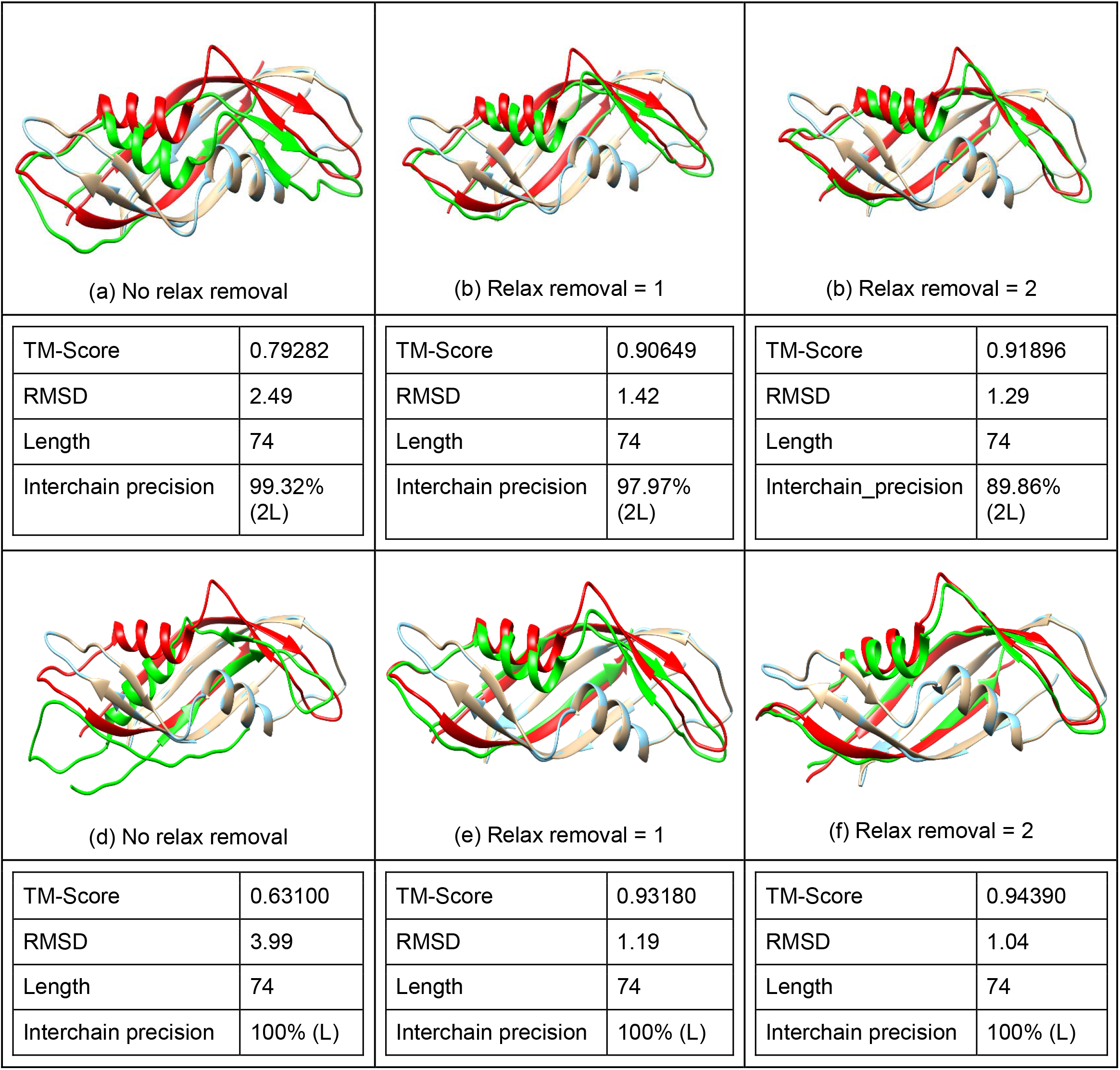
Comparison for target 1IHR between its true homodimer structure (Golden: original chain A; red: original chain B) and the structure derived from predicted contacts (Cyan: reconstructed chain A; green: reconstructed chain B). The TM-score and RMSDs were obtained using TM-Align. From (a) to (c), as we perform relax removal, the TM-score increases while RMSD decreases. But interchain precision (Top-2L) decreases slightly. Structures (d) to (f) are based on Top-L contacts, all of which have 100% precision leading to much higher TM-score and lower RMSD values. The TM-score of (d) was low (even less than (a)) due to the low overlap of the alpha-helix and some noodle regions.

## Conclusion, Limitation, and Future Work

This study demonstrates that DNCON2, a deep learning-based intrachain contact predictor, can successfully be used to predict interchain residue-residue contacts for homodimeric and homomultimeric protein complexes from the multiple sequence alignment of a monomer. Although the precision of the predictor is not high on average due to it being trained mainly for intrachain contact prediction, the prediction accuracy is still much higher than a random predictor. In some cases, our approach predicted more interchain contacts than intrachain contacts with very high precision. The results provide good evidence that deep learning tools can be used to train for such a task using co-evolutionary features obtained directly from homology-based multiple sequence alignment of a monomer. Unlike the previous works of (Hopf et al., 2014; Ovchinnikov et al., 2014; Zeng et al., 2018; Zhou et al., 2018) that requires hard-to-obtain MSAs of interlogs as input, our approach can be readily applied to any homodimers and homomultimers. Moreover, the quality of the interchain contacts directly influences the construction of the final 3D complex structure, and removing false-positive contacts through relax removal can significantly increase the TM-score of the constructed 3D structures in some cases. Our results conclude that relax removal = 2 and relaxation = 2 gives us the best precision, especially for Top-L/10 contacts.

Since DNCON2 has been trained for intrachain protein contact prediction, it has not reached the best prediction of predicting interchain contacts. Thus, our results encourage us to train a complex deep learning architecture specific to predicting interchain contacts of homodimers and homomultimers in the future. Additional co-evolutionary features like covariance matrix and pseudo-likelihood maximization (PLM) matrix may be used as input to train interchain contact predictors.

## Supporting information

Supplementary Section

## Acknowledgment

Research reported in this publication was supported in part by two NSF grants (DBI 1759934 and IIS1763246) to JC, an NIH grant (R01GM093123), and a Department of Energy grant (DE-AR0001213) to JC.

