## Supplementary Section for "Predicting interchain contacts for homodimeric and homomultimeric protein complexes using multiple sequence alignments of monomers and deep learning"

### 1. DNCON2 Deep Learning Network Architecture for Predicting Intra-chain contacts

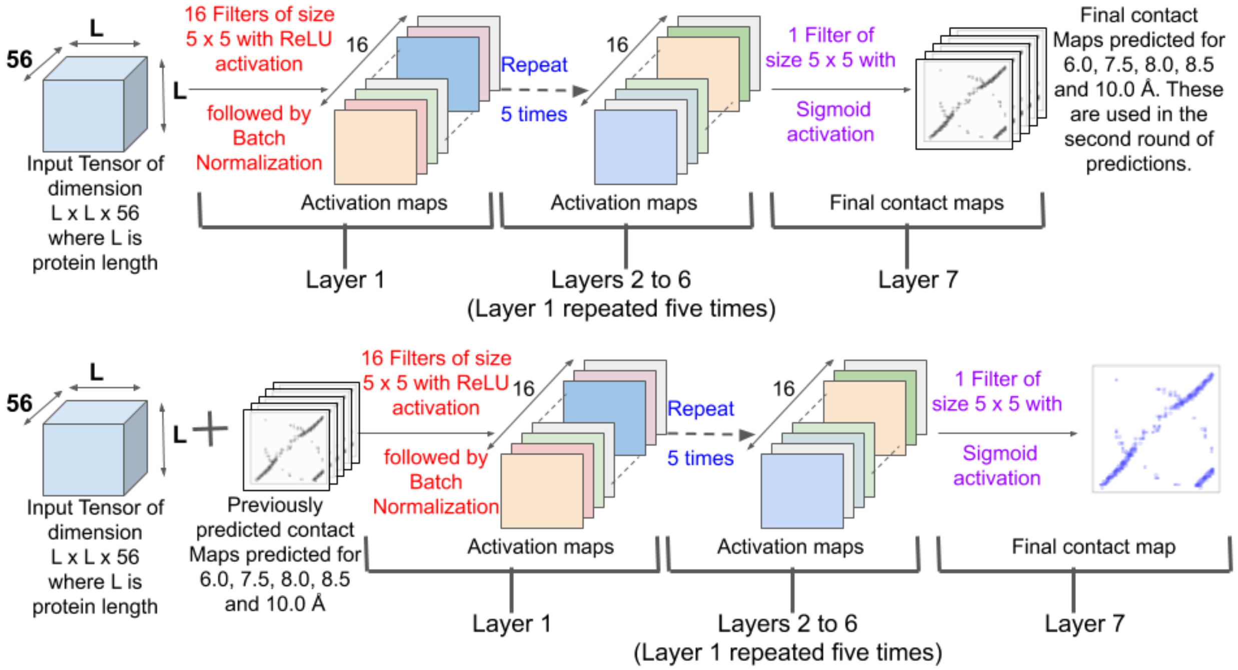


**Figure S1:** The 2D deep convolutional neural network (CNN) architecture of DNCON2. The input is an  L x L x 56 tensor (L: length of the protein), and 56 is the number of channels. (Top) The input is passed through a CNN where the first six layers consist of 16 different 5 x 5 filters, activated using the Rectified Linear Unit (ReLU) followed by batch normalization. The final output layer consists of one 5 x 5 filter, followed by sigmoid activation to predict the L x L contact map. Five different CNNs of similar architecture were trained to predict contacts at thresholds 6.0, 7.5, 8.0, 8.5, and 10.0 Å. (Bottom). These contact maps are then concatenated to the initial input tensor and then trained using a similar network to predict the final contact map at 8.0 Å. The loss function used is binary cross-entropy with Nesterov Adam (nadam) optimizer.

### 2. DNCON2 features

**Table S1.** List of features used by DNCON2. Shannon entropy, CCMPred, FreeContact, PSICOV, Mean Contact Potential, Normalized Mutual Information, and Mutual information are co-evolution-based features derived from multiple sequence alignment (MSA).

| Feature | Dimensions | Features | Channels after augmentation |
|---|---|---|---|
| Log of sequence length | Scalar | 1 | 1 |
| Log of number of sequences in the alignment | Scalar | 1 | 1 |
| Log of the number of effective number of sequences in the alignment | Scalar | 1 | 1 |
| The ratio of number of 'buried' residues and length of the protein | 1D | 1 | 1 |
| The ratio of number of 'beta-strand' residues and length of the protein | 1D | 1 | 1 |
| Ratio of number of 'helical' residues and length of the protein | 1D | 1 | 1 |
| Atchley factors normalized by sigmoid function | 1D | 5 | 10 |
| Binary predictions for helix, coil, and strand residues by SCRATCH | 1D | 3 | 6 |

| | | | |
|---|---|---|---|
| Solvent accessibility predicted by SCRATCH | 1D | 1 | 2 |
| Position Specific Scoring Matrix (PSSM) | 1D | 1 | 2 |
| PSSM Sums (divided by 100) | 1D | 1 | 2 |
| PSSM sum cosines | 1D | 1 | 2 |
| Ratio of sequence separation and length of protein | 2D | 1 | 1 |
| Flag for sequence separation between 23 and 28 | 2D | 1 | 1 |
| Flag for sequence separation between 28 and 38 | 2D | 1 | 1 |
| Flag for sequence separation between 38 and 48 | 2D | 1 | 1 |
| Flag for sequence separation 48+ | 2D | 1 | 1 |
| Probabilities of PSIPRED predictions for helix, coil, and strand residues | 1D | 3 | 6 |
| Probabilities of PSISOLV predictions for solvent accessibility | 1D | 1 | 2 |
| Pre-computed statistical potentials | 2D | 6 | 6 |
| Shannon entropy sum of the alignment columns | 2D | 1 | 1 |
| CCMpred co-evolution prediction from MSA | 2D | 1 | 1 |
| FreeContact co-evolution prediction from MSA | 2D | 1 | 1 |
| PSICOV co-evolution prediction from MSA | 2D | 1 | 1 |
| Mean contact potential from MSA | 2D | 1 | 1 |
| Normalized mutual information from MSA | 2D | 1 | 1 |
| Mutual information from MSA | 2D | 1 | 1 |
| Total | | 40 | 56 |

#### 3. Table for Contact Density distribution of proteins

**Table S2:** Table showing the contact density distribution of homodimers and homomultimers.

| | Number of proteins | |
|---|---|---|
| **Contact Density Range** | **Homodimers** | **Homomultimers** |
| 0.00-0.25 | 1291 | 1480 |
| 0.25-0.50 | 2591 | 2053 |
| 0.50-0.75 | 2085 | 1616 |
| 0.75-1.00 | 1302 | 871 |
| 1.00-1.25 | 561 | 392 |
| 1.25-1.50 | 341 | 179 |
| 1.50-1.75 | 163 | 77 |
| 1.75-2.00 | 130 | 37 |
| 2.00-2.25 | 80 | 22 |
| 2.25-2.50 | 43 | 16 |
| 2.50-2.75 | 35 | 15 |
| 2.75-3.00 | 24 | 2 |
| 3.00-3.25 | 16 | 1 |
| 3.25-3.50 | 4 | 1 |
| 3.50-3.75 | 4 | 1 |
| 3.75-4.00 | 2 | 0 |
| 4.00-4.25 | 5 | 0 |
| 4.25-4.50 | 3 | 1 |
| 4.50-4.75 | 1 | 0 |

**4. Random Prediction Precision**

**Table S3:** Table showing the precision of the random prediction of homodimers for different relaxation removal and relaxation values.

| Precision (%) | | | | | | | | |
|---|---|---|---|---|---|---|---|---|
| Relax Removal | Relaxation | Top-5 | Top-10 | Top-L/10 | Top-L/5 | Top-L/2 | Top-L | Top-2L |
| 0 | 0 | 0.69 | 0.72 | 0.69 | 0.69 | 0.68 | 0.67 | 0.65 |
| 0 | 1 | 3.07 | 3.13 | 3.09 | 3.10 | 3.09 | 3.11 | 3.04 |
| 0 | 2 | 5.73 | 5.93 | 5.84 | 5.89 | 5.83 | 5.79 | 5.61 |
| 1 | 0 | 0.66 | 0.71 | 0.70 | 0.70 | 0.69 | 0.67 | 0.59 |
| 1 | 1 | 2.95 | 3.01 | 2.98 | 3.04 | 3.04 | 3.03 | 2.69 |
| 1 | 2 | 5.57 | 5.74 | 5.66 | 5.76 | 5.71 | 5.64 | 4.99 |
| 2 | 0 | 0.70 | 0.73 | 0.70 | 0.73 | 0.69 | 0.68 | 0.54 |
| 2 | 1 | 2.95 | 3.01 | 2.95 | 3.01 | 3.02 | 3.01 | 2.43 |
| 2 | 2 | 5.54 | 5.67 | 5.55 | 5.65 | 5.56 | 5.51 | 4.44 |

**Table S4:** Table showing the precision of the random prediction of homomultimers for different relaxation removal and relaxation values.

| Precision (%) | | | | | | | | |
|---|---|---|---|---|---|---|---|---|
| Relax Removal | Relaxation | Top-5 | Top-10 | Top-L/10 | Top-L/5 | Top-L/2 | Top-L | Top-2L |
| 0 | 0 | 0.52 | 0.56 | 0.59 | 0.61 | 0.61 | 0.60 | 0.58 |
| 0 | 1 | 2.67 | 2.74 | 2.77 | 2.76 | 2.74 | 2.70 | 2.63 |
| 0 | 2 | 4.92 | 5.07 | 5.08 | 5.12 | 5.11 | 5.10 | 4.94 |
| 1 | 0 | 0.52 | 0.56 | 0.58 | 0.59 | 0.60 | 0.59 | 0.52 |
| 1 | 1 | 2.59 | 2.66 | 2.68 | 2.70 | 2.66 | 2.65 | 2.31 |
| 1 | 2 | 4.92 | 4.98 | 4.96 | 4.98 | 4.96 | 4.97 | 4.33 |
| 2 | 0 | 0.49 | 0.54 | 0.56 | 0.57 | 0.59 | 0.59 | 0.46 |
| 2 | 1 | 2.52 | 2.60 | 2.63 | 2.63 | 2.61 | 2.61 | 2.05 |
| 2 | 2 | 4.73 | 4.83 | 4.86 | 4.82 | 4.84 | 4.83 | 3.79 |
